## Supplementary Files 1-12 for "Neurophysiological trajectories in Alzheimer’s disease progression"

**Supplementary File 1:**
**Demographics and Neuropsychological assessments**.

| Variable | Controls ($n = 70$) | Patients with AD ($n = 78$) |
|---|---|---|
| Age (years) | $70.5 \pm 8.28$ | $63.9 \pm 8.93$ |
| Sex (% female) | 41 (59%) | 50 (64%) |
| Handedness (% right) | 54 (77%) | 65 (83%) |
| Education (years) | $17.43 \pm 1.97$ | $16.6 \pm 2.47$ |
| MMSE | $29.36 \pm 1.00$ | $22.67 \pm 4.79$ |
| CDR | $0.00 \pm 0.00$ | $0.83 \pm 0.46$ |
| CDR-SOB | $0.03 \pm 0.12$ | $4.32 \pm 3.13$ |
| Modified trails | $0.65 \pm 0.25$ | $0.24 \pm 0.21$ |
| Design fluency | $12.82 \pm 3.56$ | $6.33 \pm 3.74$ |
| Phonemic fluency | $17.52 \pm 5.40$ | $10.75 \pm 5.14$ |
| Category fluency | $23.93 \pm 4.88$ | $12.14 \pm 6.29$ |
| Digit span forwards | $7.16 \pm 1.34$ | $5.04 \pm 1.52$ |
| Digit span backwards | $5.75 \pm 1.32$ | $3.51 \pm 1.30$ |
| Processing speed | $86.72 \pm 15.51$ | $52.09 \pm 24.08$ |
| Stroop inhibition | $52.06 \pm 14.07$ | $23.59 \pm 16.03$ |
| Modified Rey copy | $15.50 \pm 0.77$ | $11.55 \pm 5.01$ |
| VOSP number location | $9.13 \pm 1.23$ | $6.97 \pm 2.70$ |
| Repetition | $4.83 \pm 0.42$ | $3.52 \pm 1.41$ |
| CVLT learning (%) | $66.31 \pm 13.34$ | $38.93 \pm 14.69$ |
| CVLT short delay recall (%) | $74.53 \pm 17.89$ | $43.85 \pm 27.33$ |
| CVLT long delay recall (%) | $77.84 \pm 18.05$ | $27.56 \pm 31.07$ |
| Modified Rey recall | $12.12 \pm 2.79$ | $5.03 \pm 4.36$ |
| Geriatric Depression Scale | $2.62 \pm 2.58$ | $6.85 \pm 4.46$ |

- Mini Mental State Examination (MMSE) (score out of 30).
- Clinical Dementia Rating (CDR) (score out of 0, 0.5, 1, 2, 3).
- CDR sum of boxes (CDR-SOB) (score out of 18).
- Modified trails assess set-shifting and are indicated by the number of correct lines drawn within 60 seconds in the modified trail making test, which requires the subject to serially alternate between numbers and days of the week.
- Design fluency, assessed using the filled-dots condition from the design fluency subscale of the Delis-Kaplan Executive Function Scale (DKEFS), was scored as the number of correct designs generated within 60 seconds.
- Category fluency indicates the number of animals listed within 60 seconds.
- Phonemic fluency indicates the number of words starting from the letter 'D' listed within 60 seconds.
- Digit span forwards assess auditory attention and are indicated by the number of digits correctly repeated in the same order from a list read by the examiner.
- Digit span backwards assess the verbal working memory and are indicated by the number of digits correctly repeated backwards from a list read by the examiner.
- Processing speed is the number of words correctly read in the congruent Stroop test within 60 seconds.
- Stroop inhibition is the number of words correctly read in the incongruent Stroop test within 60 seconds.
- Modified Rey copy assesses visual construction copy of the Benson figure (score out of 17).
- VOSP number location is assessed by the number location task of the Visual Object and Space Perception Battery (score out of 10).
- Repetition is assessed by participants repeating five phonemically complex sentences following the examiner (score out of 5).
- CVLT leaning indicates the cumulative number of words recalled from trails 1-5 in 16 item CVLT word list (score out of 80) for controls and 9 item CVLT word list (total out of 45) for AD.
- CVLT short delay recall indicates the number of words recalled after a 30 second delay from 16 item CVLT word list (score out of 16) for controls and from 9 item CVLT word list (score out of 9 for AD).
- CVLT long delay recall indicates the number of words recalled after a 10-minute delay from 16 item CVLT word list (score out of 16) for controls and from 9 item CVLT word list (score out of 9 for AD).
- Modified Ray recall is construction of Benson figure from memory after 10 minutes (score out of 17).
- Geriatric Depression Scale (GDS) (score out of 30).

**Supplementary File 2:**
**The 94 cortical/subcortical anatomical regions included in the AAL3 atlas**.

| No. (left, right) | Anatomical description | Abbreviation |
|---|---|---|
| 1, 2 | Precentral gyrus | PreCG |
| 3, 4 | Superior frontal gyrus-dorsolateral | SFG |
| 5, 6 | Middle frontal gyrus | MFG |
| 7, 8 | Inferior frontal gyrus-opercular part | IFGoper |
| 9, 10 | Inferior frontal gyrus-triangular part | IFGtria |
| 11, 12 | IFG pars orbitalis | IFGorb |
| 13, 14 | Rolandic operculum | ROL |
| 15, 16 | Supplementary motor area | SMA |
| 17, 18 | Olfactory cortex | OLF |
| 19, 20 | Superior frontal gyrus-medial | SFGmed |
| 21, 22 | Superior frontal gyrus-medial orbital | PFCvent |
| 23, 24 | Gyrus rectus | REC |
| 25, 26 | Medial orbital gyrus | OFCmed |
| 27, 28 | Anterior orbital gyrus | OFCant |
| 29, 30 | Posterior orbital gyrus | OFCpost |
| 31, 32 | Lateral orbital gyrus | OFClat |
| 33, 34 | Insula | INS |
| 35, 36 | Anterior cingulate & paracingulate gyri | ACC |
| 37, 38 | Middle cingulate & paracingulate gyri | MCC |
| 39, 40 | Posterior cingulate gyrus | PCC |
| 41, 42 | Hippocampus | HIP |
| 43, 44 | Parahippocampal gyrus | PHG |
| 45, 46 | Amygdala | AMYG |
| 47, 48 | Calcarine fissure and surrounding cortex | CAL |
| 49, 50 | Cuneus | CUN |
| 51, 52 | Lingual gyrus | LING |
| 53, 54 | Superior occipital gyrus | SOG |
| 55, 56 | Middle occipital gyrus | MOG |
| 57, 58 | Inferior occipital gyrus | IOG |
| 59, 60 | Fusiform gyrus | FFG |
| 61, 62 | Postcentral gyrus | PoCG |
| 63, 64 | Superior parietal gyrus | SPG |
| 65, 66 | Inferior parietal gyrus | IPG |
| 67, 68 | SupraMarginal gyrus | SMG |
| 69, 70 | Angular gyrus | ANG |
| 71, 72 | Precuneus | PCUN |
| 73, 74 | Paracentral lobule | PCL |
| 75, 76 | Caudate nucleus | CAU |
| 77, 78 | Lenticular nucleus-Putamen | PUT |
| 79, 80 | Lenticular nucleus-Pallidum | PAL |
| 81, 82 | Thalamus | THA |
| 83, 84 | Heschls gyrus | HES |
| 85, 86 | Superior temporal gyrus | STG |
| 87, 88 | Temporal pole: superior temporal gyrus | TPOsup |
| 89, 90 | Middle temporal gyrus | MTG |
| 91, 92 | Temporal pole: middle temporal gyrus | TPOmid |
| 93, 94 | Inferior temporal gyrus | ITG |

### Supplementary File 3:

**Direct evaluation of the data likelihoods for all possible $z$-score event sequences in the AC-EBM.** In this example, we select the optimal set of $z$-score events; $z_{\mathrm{PHG}}$ = [E1 E2 E3] = [0.0964 0.4708 2.4519] and $z_{\mathrm{MMSE}}$ = [E4 E5 E6] = [0.5109 4.4116 6.2417] (see Step 2 in Figure 1—figure supplement 1). In an AC-EBM, there are total 20 ordered arrangements of the $z$-score events. This evaluation indicates that the data likelihood, $\log P(Z|S)$, is maximized when $S$ =[E1 E2 E4 E5 E6 E3] (labeled $*$ in the table), as shown in Step 2 in Figure 1—figure supplement 1. The second highest likelihood was obtained when $S$ =[E1 E4 E2 E5 E6 E3] (labeled $**$ in the table). It is worth noting that $P_{**}/P_{*}$ ~ 2.66E-07. This indicates that it is extremely rare for sequences other than the most likely sequence ($S$ =[E1 E2 E4 E5 E6 E3]) to occur in the MCMC sampling. This is the reason why we obtained the positional variance diagram without positional uncertainty for the optimal set of $z$-score events in the AC-EBM.

| | Sequence $S$ | $\log P(Z|S)$ |
|---|---|---|
| | E1 E2 E3 E4 E5 E6 | -473.08 |
| | E1 E2 E4 E3 E5 E6 | -435.16 |
| | E1 E2 E4 E5 E3 E6 | -407.33 |
| $*$ | E1 E2 E4 E5 E6 E3 | -353.69 |
| | E1 E4 E2 E3 E5 E6 | -430.07 |
| | E1 E4 E2 E5 E3 E6 | -417.75 |
| $**$ | E1 E4 E2 E5 E6 E3 | -368.83 |
| | E1 E4 E5 E2 E3 E6 | -434.35 |
| | E1 E4 E5 E2 E6 E3 | -396.72 |
| | E1 E4 E5 E6 E2 E3 | -423.01 |
| | E4 E1 E2 E3 E5 E6 | -462.35 |
| | E4 E1 E2 E5 E3 E6 | -461.09 |
| | E4 E1 E2 E5 E6 E3 | -412.08 |
| | E4 E1 E5 E2 E3 E6 | -487.32 |
| | E4 E1 E5 E2 E6 E3 | -449.11 |
| | E4 E1 E5 E6 E2 E3 | -473.24 |
| | E4 E5 E1 E2 E3 E6 | -545.23 |
| | E4 E5 E1 E2 E6 E3 | -512.36 |
| | E4 E5 E1 E6 E2 E3 | -540.27 |
| | E4 E5 E6 E1 E2 E3 | -596.36 |

### Supplementary File 4:

**Top 10 regions with significant group differences in GM volumes comparison between AD patients and controls.** Negative $T$-value represents that a mean regional volume in the AD group is smaller than that in controls. The degree of freedom $df = 143$. A value of 0.000E+00 denotes 2.2204E-16 (double precision).

| Regions (AAL3 atlas) | $T$-value | $p$-value | $q$-value |
|---|---|---|---|
| Left Inferior temporal gyrus | -10.649 | 0.000E+00 | 0.000E+00 |
| Left Middle temporal gyrus | -10.605 | 0.000E+00 | 0.000E+00 |
| Right Middle temporal gyrus | -10.253 | 0.000E+00 | 0.000E+00 |
| Left Fusiform gyrus | -9.758 | 0.000E+00 | 0.000E+00 |
| Left Hippocampus | -9.457 | 0.000E+00 | 0.000E+00 |
| Left Parahippocampal gyrus | -8.808 | 3.997E-15 | 6.262E-14 |
| Right Inferior temporal gyrus | -8.614 | 1.177E-14 | 1.580E-13 |
| Left Precuneus | -8.585 | 1.399E-14 | 1.644E-13 |
| Right Parahippocampal gyrus | -8.524 | 1.976E-14 | 2.064E-13 |
| Right Hippocampus | -8.411 | 3.775E-14 | 3.548E-13 |

### Supplementary File 5:

**Top 10 regions with significant group differences in long-range synchrony between patients with AD and controls.** Negative $T$-value represents that a mean regional metric in patients with AD is smaller than that in controls. The degree of freedom $df = 145$.

| Frequency band | Regions (AAL3 atlas) | $T$-value | $p$-value | $q$-value |
|---|---|---|---|---|
| delta-theta | Left Precentral gyrus | 3.827 | 1.925E-04 | 9.938E-03 |
| | Right Superior frontal gyrus-dorsolateral | 3.624 | 4.011E-04 | 9.938E-03 |
| | Right Superior frontal gyrus-medial | 3.521 | 5.741E-04 | 9.938E-03 |
| | Right Anterior cingulate & paracingulate gyri | 3.518 | 5.802E-04 | 9.938E-03 |
| | Left Superior frontal gyrus-medial | 3.509 | 5.993E-04 | 9.938E-03 |
| | Right Middle frontal gyrus | 3.493 | 6.343E-04 | 9.938E-03 |
| | Right Supplementary motor area | 3.425 | 8.011E-04 | 1.076E-02 |
| | Left Middle frontal gyrus | 3.372 | 9.585E-04 | 1.092E-02 |
| | Left Superior frontal gyrus-dorsolateral | 3.346 | 1.046E-03 | 1.092E-02 |
| | Left Inferior frontal gyrus-opercular part | 3.310 | 1.178E-03 | 1.107E-02 |
| alpha | Left SupraMarginal gyrus | -7.638 | 2.735E-12 | 2.571E-10 |
| | Left Rolandic operculum | -6.751 | 3.280E-10 | 1.184E-08 |
| | Left Middle temporal gyrus | -6.665 | 5.152E-10 | 1.184E-08 |
| | Right Fusiform gyrus | -6.651 | 5.526E-10 | 1.184E-08 |
| | Left Superior temporal gyrus | -6.626 | 6.299E-10 | 1.184E-08 |
| | Left Fusiform gyrus | -6.469 | 1.418E-09 | 2.222E-08 |
| | Left Heschls gyrus | -6.391 | 2.117E-09 | 2.843E-08 |
| | Left Inferior parietal gyrus | -6.196 | 5.679E-09 | 5.959E-08 |
| | Right Hippocampus | -6.192 | 5.800E-09 | 5.959E-08 |
| | Left Thalamus | -6.174 | 6.339E-09 | 5.959E-08 |
| beta | Right Middle temporal gyrus | -8.237 | 9.459E-14 | 8.892E-12 |
| | Right Angular gyrus | -7.553 | 4.387E-12 | 2.062E-10 |
| | Left Middle temporal gyrus | -7.455 | 7.521E-12 | 2.301E-10 |
| | Left Inferior temporal gyrus | -7.406 | 9.793E-12 | 2.301E-10 |
| | Left Middle occipital gyrus | -7.035 | 7.308E-11 | 1.160E-09 |
| | Left Angular gyrus | -7.032 | 7.404E-11 | 1.160E-09 |
| | Left Superior temporal gyrus | -6.817 | 2.326E-10 | 3.124E-09 |
| | Right Fusiform gyrus | -6.647 | 5.667E-10 | 6.659E-09 |
| | Right Inferior temporal gyrus | -6.383 | 2.208E-09 | 2.082E-08 |
| | Right Inferior parietal gyrus | -6.382 | 2.215E-09 | 2.082E-08 |

### Supplementary File 6:

**Top 10 regions with significant group differences in local synchrony between patients with AD and controls.** Negative $T$-value represents that a mean regional metric in patients with AD is smaller than that in controls. The degree of freedom $df = 145$. A value of 0.000E+00 denotes 2.2204E-16 (double precision).

| Frequency band | Regions (AAL3 atlas) | $T$-value | $p$-value | $q$-value |
|---|---|---|---|---|
| delta-theta | Left Inferior temporal gyrus | 11.235 | 0.000E+00 | 0.000E+00 |
| | Right Fusiform gyrus | 10.937 | 0.000E+00 | 0.000E+00 |
| | Right Inferior occipital gyrus | 10.655 | 0.000E+00 | 0.000E+00 |
| | Left Middle temporal gyrus | 10.643 | 0.000E+00 | 0.000E+00 |
| | Left Superior temporal gyrus | 10.616 | 0.000E+00 | 0.000E+00 |
| | Left Parahippocampal gyrus | 10.607 | 0.000E+00 | 0.000E+00 |
| | Left Heschls gyrus | 10.534 | 0.000E+00 | 0.000E+00 |
| | Left Rolandic operculum | 10.521 | 0.000E+00 | 0.000E+00 |
| | Left Inferior occipital gyrus | 10.516 | 0.000E+00 | 0.000E+00 |
| | Left Fusiform gyrus | 10.506 | 0.000E+00 | 0.000E+00 |
| alpha | Left Inferior temporal gyrus | -7.805 | 1.082E-12 | 1.017E-10 |
| | Left Fusiform gyrus | -7.529 | 4.994E-12 | 2.347E-10 |
| | Left Parahippocampal gyrus | -7.327 | 1.512E-11 | 4.058E-10 |
| | Left Inferior occipital gyrus | -7.302 | 1.727E-11 | 4.058E-10 |
| | Right Amygdala | -6.793 | 2.632E-10 | 4.949E-09 |
| | Right Fusiform gyrus | -6.626 | 6.291E-10 | 9.856E-09 |
| | Right Temporal pole: superior temporal gyrus | -6.562 | 8.771E-10 | 1.156E-08 |
| | Right Inferior occipital gyrus | -6.522 | 1.080E-09 | 1.156E-08 |
| | Right Parahippocampal gyrus | -6.517 | 1.107E-09 | 1.156E-08 |
| | Left Middle temporal gyrus | -6.485 | 1.305E-09 | 1.227E-08 |
| beta | Left Angular gyrus | -8.008 | 3.468E-13 | 3.260E-11 |
| | Left Superior temporal gyrus | -7.747 | 1.497E-12 | 7.037E-11 |
| | Right Angular gyrus | -7.416 | 9.314E-12 | 2.918E-10 |
| | Right Hippocampus | -7.272 | 2.041E-11 | 4.795E-10 |
| | Left Heschls gyrus | -7.194 | 3.114E-11 | 5.305E-10 |
| | Left Lenticular nucleus-Pallidum | -7.178 | 3.386E-11 | 5.305E-10 |
| | Left Hippocampus | -7.089 | 5.453E-11 | 7.299E-10 |
| | Right Middle occipital gyrus | -7.065 | 6.212E-11 | 7.299E-10 |
| | Right Heschls gyrus | -6.998 | 8.913E-11 | 9.309E-10 |
| | Left Parahippocampal gyrus | -6.965 | 1.058E-10 | 9.944E-10 |

**Supplementary File 7:**

**Top $10$ regions with significant weighted mean differences ($q < 0.05$, FDR corrected) in long-range synchrony between stages 4 and 1 (Figure 2E in the main text)**. The $p$- and $q$-values of 0.000E+00 denote a value less than $1/50,000$, where $50,000$ is the number of the bootstrap samplings in the non-parametric tests.

| Frequency band | Regions (AAL3 atlas) | $\delta z$ [Stages 4 vs.1] | $p$-value | $q$-value |
|---|---|---|---|---|
| alpha | Left SupraMarginal gyrus | -0.975 | 2.000E-05 | 3.760E-04 |
| | Left Rolandic operculum | -0.933 | 8.000E-05 | 9.400E-04 |
| | Left Fusiform gyrus | -0.911 | 2.000E-05 | 3.760E-04 |
| | Left Thalamus | -0.895 | 2.000E-05 | 3.760E-04 |
| | Right Temporal pole: middle temporal gyrus | -0.878 | 2.000E-05 | 3.760E-04 |
| | Left Heschls gyrus | -0.856 | 4.000E-05 | 6.267E-04 |
| | Left Lenticular nucleus-Pallidum | -0.839 | 2.000E-04 | 1.216E-03 |
| | Left Lenticular nucleus-Putamen | -0.836 | 1.200E-04 | 1.128E-03 |
| | Right Inferior temporal gyrus | -0.834 | 3.400E-04 | 1.522E-03 |
| | Left Inferior parietal gyrus | -0.833 | 2.000E-04 | 1.216E-03 |
| beta | Right Middle temporal gyrus | -1.160 | 0.000E+00 | 0.000E+00 |
| | Left Inferior temporal gyrus | -1.016 | 0.000E+00 | 0.000E+00 |
| | Left Angular gyrus | -1.015 | 0.000E+00 | 0.000E+00 |
| | Left Superior temporal gyrus | -0.972 | 2.000E-05 | 2.089E-04 |
| | Right Inferior temporal gyrus | -0.935 | 0.000E+00 | 0.000E+00 |
| | Right Fusiform gyrus | -0.930 | 2.000E-05 | 2.089E-04 |
| | Left Middle temporal gyrus | -0.925 | 4.000E-05 | 2.892E-04 |
| | Right Superior temporal gyrus | -0.924 | 6.000E-05 | 3.760E-04 |
| | Right Inferior parietal gyrus | -0.921 | 4.000E-05 | 2.892E-04 |
| | Right Angular gyrus | -0.917 | 1.600E-04 | 5.013E-04 |

### Supplementary File 8:

**Top $10$ regions with significant weighted mean differences ($q < 0.05$, FDR corrected) in local synchrony between stages 4 and 1 (Figure 2F in the main text)**. The $p$-and $q$-values of 0.000E+00 denote a value less than $1/50,000$, where $50,000$ is the number of the bootstrap samplings in the non-parametric tests.

| Frequency band | Regions (AAL3 atlas) | $\delta z$ | $p$-value | $q$-value |
|---|---|---|---|---|
| delta-theta | Right Middle occipital gyrus | 2.363 | 2.400E-04 | 1.074E-03 |
| | Left Middle temporal gyrus | 2.275 | 0.000E+00 | 0.000E+00 |
| | Left SupraMarginal gyrus | 2.272 | 6.000E-05 | 8.057E-04 |
| | Left Angular gyrus | 2.218 | 1.200E-04 | 8.847E-04 |
| | Left Rolandic operculum | 2.167 | 6.000E-05 | 8.057E-04 |
| | Left Superior temporal gyrus | 2.160 | 6.000E-05 | 8.057E-04 |
| | Left Middle occipital gyrus | 2.081 | 1.400E-04 | 8.847E-04 |
| | Right Superior occipital gyrus | 2.073 | 6.000E-04 | 1.312E-03 |
| | Left Superior occipital gyrus | 2.064 | 8.000E-05 | 8.356E-04 |
| | Left Heschls gyrus | 2.061 | 8.000E-05 | 8.356E-04 |
| alpha | Left Fusiform gyrus | -1.170 | 0.000E+00 | 0.000E+00 |
| | Left Inferior temporal gyrus | -1.051 | 2.200E-04 | 1.034E-02 |
| | Left Parahippocampal gyrus | -0.961 | 5.600E-04 | 1.065E-02 |
| | Left Inferior occipital gyrus | -0.933 | 8.600E-04 | 1.065E-02 |
| | Left Hippocampus | -0.928 | 9.400E-04 | 1.065E-02 |
| | Left Amygdala | -0.912 | 8.800E-04 | 1.065E-02 |
| | Right Fusiform gyrus | -0.891 | 1.340E-03 | 1.065E-02 |
| | Right Temporal pole: superior temporal gyrus | -0.862 | 7.000E-04 | 1.065E-02 |
| | Left Middle temporal gyrus | -0.835 | 2.080E-03 | 1.450E-02 |
| | Left Lingual gyrus | -0.827 | 7.800E-04 | 1.065E-02 |
| beta | Left Superior temporal gyrus | -1.132 | 2.800E-04 | 7.708E-03 |
| | Left Heschls gyrus | -1.076 | 1.600E-04 | 7.520E-03 |
| | Left Angular gyrus | -1.048 | 5.000E-04 | 7.708E-03 |
| | Left Rolandic operculum | -1.026 | 4.400E-04 | 7.708E-03 |
| | Left Parahippocampal gyrus | -1.011 | 4.000E-05 | 3.760E-03 |
| | Left Insula | -0.994 | 5.600E-04 | 7.708E-03 |
| | Right Heschls gyrus | -0.953 | 8.200E-04 | 7.708E-03 |
| | Left SupraMarginal gyrus | -0.951 | 7.600E-04 | 7.708E-03 |
| | Left Lenticular nucleus-Putamen | -0.934 | 6.400E-04 | 7.708E-03 |
| | Left Inferior temporal gyrus | -0.919 | 8.200E-04 | 7.708E-03 |

**Supplementary File 9:**

**Pairs of stages with significant weighted-mean differences ($q$<0.05, FDR corrected) in the trajectories of delta-theta-, alpha-, and beta-band long-range synchrony in the SAC-EBMs (Figure 3B, F, J in the main text).** The $p$-values of 0.000E+00 denote a value less than $1/50,000$, where $50,000$ is the number of bootstrap samplings.

**delta-theta**

| Stages | $p$-value | $q$-value |
|---|---|---|
| (4,1) | 5.560E-03 | 1.787E-02 |
| (5,1) | 4.180E-03 | 1.710E-02 |
| (6,1) | 0.000E+00 | 0.000E+00 |
| (6,2) | 1.200E-04 | 1.200E-03 |
| (6,3) | 5.020E-03 | 1.738E-02 |
| (6,4) | 1.258E-02 | 2.979E-02 |
| (7,1) | 0.000E+00 | 0.000E+00 |
| (7,2) | 1.600E-04 | 1.200E-03 |
| (7,3) | 3.620E-03 | 1.629E-02 |
| (7,4) | 9.040E-03 | 2.542E-02 |
| (8,1) | 0.000E+00 | 0.000E+00 |
| (8,2) | 9.200E-04 | 5.400E-03 |
| (8,3) | 1.250E-02 | 2.979E-02 |
| (9,1) | 1.080E-03 | 5.400E-03 |
| (9,2) | 4.920E-03 | 1.738E-02 |
| (10,1) | 1.400E-04 | 1.200E-03 |
| (10,2) | 1.040E-03 | 5.400E-03 |
| (10,3) | 7.100E-03 | 2.130E-02 |
| (10,4) | 1.226E-02 | 2.979E-02 |

**alpha**

| Stages | $p$-value | $q$-value |
|---|---|---|
| (2,1) | 4.000E-05 | 1.800E-04 |
| (3,1) | 0.000E+00 | 0.000E+00 |
| (4,1) | 0.000E+00 | 0.000E+00 |
| (4,2) | 2.320E-03 | 7.457E-03 |
| (5,1) | 0.000E+00 | 0.000E+00 |
| (5,2) | 4.000E-05 | 1.800E-04 |
| (6,1) | 0.000E+00 | 0.000E+00 |
| (6,2) | 1.000E-04 | 4.091E-04 |
| (7,1) | 0.000E+00 | 0.000E+00 |
| (7,2) | 1.200E-04 | 4.500E-04 |
| (8,1) | 0.000E+00 | 0.000E+00 |
| (8,2) | 1.980E-03 | 6.854E-03 |
| (9,1) | 0.000E+00 | 0.000E+00 |
| (9,2) | 8.940E-03 | 2.514E-02 |
| (10,1) | 0.000E+00 | 0.000E+00 |
| (10,2) | 2.660E-03 | 7.980E-03 |

**beta**

| Stages | $p$-value | $q$-value |
|---|---|---|
| (2,1) | 0.000E+00 | 0.000E+00 |
| (3,1) | 0.000E+00 | 0.000E+00 |
| (4,1) | 0.000E+00 | 0.000E+00 |
| (4,2) | 8.560E-03 | 2.568E-02 |
| (5,1) | 0.000E+00 | 0.000E+00 |
| (5,2) | 5.200E-04 | 1.950E-03 |
| (6,1) | 0.000E+00 | 0.000E+00 |
| (6,2) | 6.000E-04 | 2.077E-03 |
| (7,1) | 0.000E+00 | 0.000E+00 |
| (7,2) | 2.000E-05 | 9.000E-05 |
| (7,3) | 1.390E-02 | 3.909E-02 |
| (8,1) | 0.000E+00 | 0.000E+00 |
| (8,2) | 3.400E-04 | 1.391E-03 |
| (9,1) | 2.000E-05 | 9.000E-05 |
| (10,1) | 0.000E+00 | 0.000E+00 |
| (10,2) | 2.440E-03 | 7.843E-03 |

### Supplementary File 10:

**Top $10$ regions with significant weighted-mean differences ($q < 0.05$, FDR corrected) in regional variations of long-range synchrony during preclinical stages (stages 5 vs 1) [Figure 3D, H, L in the main text].** The $p$- and $q$-values of 0.000E+00 denote a value less than $1/50,000$, where $50,000$ is the number of bootstrap samplings.

| Frequency band | Regions (AAL3 atlas) | $\delta z$ | $p$-value | $q$-value |
|---|---|---|---|---|
| alpha | Left Thalamus | -1.588 | 0.000E+00 | 0.000E+00 |
| | Left Rolandic operculum | -1.558 | 0.000E+00 | 0.000E+00 |
| | Right Olfactory cortex | -1.553 | 0.000E+00 | 0.000E+00 |
| | Left Fusiform gyrus | -1.547 | 0.000E+00 | 0.000E+00 |
| | Right Amygdala | -1.540 | 0.000E+00 | 0.000E+00 |
| | Left SupraMarginal gyrus | -1.527 | 0.000E+00 | 0.000E+00 |
| | Left Lenticular nucleus-Putamen | -1.507 | 0.000E+00 | 0.000E+00 |
| | Left Lenticular nucleus-Pallidum | -1.501 | 0.000E+00 | 0.000E+00 |
| | Right SupraMarginal gyrus | -1.481 | 0.000E+00 | 0.000E+00 |
| | Right Parahippocampal gyrus | -1.480 | 0.000E+00 | 0.000E+00 |
| beta | Left Inferior temporal gyrus | -1.771 | 0.000E+00 | 0.000E+00 |
| | Right Middle temporal gyrus | -1.711 | 0.000E+00 | 0.000E+00 |
| | Left Thalamus | -1.709 | 0.000E+00 | 0.000E+00 |
| | Left Angular gyrus | -1.707 | 0.000E+00 | 0.000E+00 |
| | Right Lenticular nucleus-Pallidum | -1.688 | 0.000E+00 | 0.000E+00 |
| | Right Superior frontal gyrus-dorsolateral | -1.676 | 0.000E+00 | 0.000E+00 |
| | Right Lenticular nucleus-Putamen | -1.662 | 0.000E+00 | 0.000E+00 |
| | Left Fusiform gyrus | -1.659 | 0.000E+00 | 0.000E+00 |
| | Left Lenticular nucleus-Pallidum | -1.647 | 0.000E+00 | 0.000E+00 |
| | Left Hippocampus | -1.644 | 0.000E+00 | 0.000E+00 |

### Supplementary File 11:

**Pairs of stages with significant weighted-mean differences ($q$<0.05, FDR corrected) in the trajectories of delta-theta-, alpha-, and beta-band local synchrony in the SAC-EBMs (Figure 4B, F, J in the main text).** The $p$- and $q$-values of 0.000E+00 denote a value less than $1/50,000$, where $50,000$ is the number of bootstrap samplings.

**delta-theta**

| Stages | $p$-value | $q$-value |
|---|---|---|
| (6,1) | 0.000E+00 | 0.000E+00 |
| (6,2) | 0.000E+00 | 0.000E+00 |
| (6,3) | 1.000E-04 | 1.957E-04 |
| (6,4) | 8.400E-04 | 1.512E-03 |
| (6,5) | 1.262E-02 | 2.103E-02 |
| (7,1) | 0.000E+00 | 0.000E+00 |
| (7,2) | 0.000E+00 | 0.000E+00 |
| (7,3) | 0.000E+00 | 0.000E+00 |
| (7,4) | 0.000E+00 | 0.000E+00 |
| (7,5) | 0.000E+00 | 0.000E+00 |
| (7,6) | 6.140E-03 | 1.063E-02 |
| (8,1) | 0.000E+00 | 0.000E+00 |
| (8,2) | 0.000E+00 | 0.000E+00 |
| (8,3) | 0.000E+00 | 0.000E+00 |
| (8,4) | 0.000E+00 | 0.000E+00 |
| (8,5) | 0.000E+00 | 0.000E+00 |
| (8,6) | 2.448E-02 | 3.934E-02 |
| (9,1) | 0.000E+00 | 0.000E+00 |
| (9,2) | 0.000E+00 | 0.000E+00 |
| (9,3) | 0.000E+00 | 0.000E+00 |
| (9,4) | 2.000E-05 | 4.091E-05 |
| (9,5) | 1.800E-04 | 3.375E-04 |
| (10,1) | 0.000E+00 | 0.000E+00 |
| (10,2) | 0.000E+00 | 0.000E+00 |
| (10,3) | 0.000E+00 | 0.000E+00 |
| (10,4) | 0.000E+00 | 0.000E+00 |
| (10,5) | 0.000E+00 | 0.000E+00 |
| (10,6) | 2.000E-05 | 4.091E-05 |

**alpha**

| Stages | $p$-value | $q$-value |
|---|---|---|
| (3,1) | 2.720E-03 | 5.322E-03 |
| (3,2) | 2.864E-02 | 4.341E-02 |
| (4,1) | 0.000E+00 | 0.000E+00 |
| (4,2) | 2.600E-04 | 5.850E-04 |
| (5,1) | 0.000E+00 | 0.000E+00 |
| (5,2) | 0.000E+00 | 0.000E+00 |
| (5,3) | 7.020E-03 | 1.215E-02 |
| (6,1) | 0.000E+00 | 0.000E+00 |
| (6,2) | 0.000E+00 | 0.000E+00 |
| (6,3) | 1.600E-04 | 3.789E-04 |
| (6,4) | 1.334E-02 | 2.144E-02 |
| (7,1) | 0.000E+00 | 0.000E+00 |
| (7,2) | 0.000E+00 | 0.000E+00 |
| (7,3) | 2.000E-05 | 5.625E-05 |
| (7,4) | 3.980E-03 | 7.463E-03 |
| (8,1) | 0.000E+00 | 0.000E+00 |
| (8,2) | 0.000E+00 | 0.000E+00 |
| (8,3) | 4.000E-05 | 1.059E-04 |
| (8,4) | 4.400E-03 | 7.920E-03 |
| (9,1) | 0.000E+00 | 0.000E+00 |
| (9,2) | 0.000E+00 | 0.000E+00 |
| (9,3) | 0.000E+00 | 0.000E+00 |
| (9,4) | 6.000E-05 | 1.500E-04 |
| (9,5) | 9.600E-04 | 2.057E-03 |
| (9,6) | 2.894E-02 | 4.341E-02 |
| (10,1) | 0.000E+00 | 0.000E+00 |
| (10,2) | 0.000E+00 | 0.000E+00 |
| (10,3) | 2.000E-05 | 5.625E-05 |
| (10,4) | 1.340E-03 | 2.741E-03 |
| (10,5) | 8.320E-03 | 1.387E-02 |

**beta**

| Stages | $p$-value | $q$-value |
|---|---|---|
| (2,1) | 1.540E-03 | 3.150E-03 |
| (3,1) | 0.000E+00 | 0.000E+00 |
| (4,1) | 0.000E+00 | 0.000E+00 |
| (4,2) | 5.360E-03 | 8.933E-03 |
| (5,1) | 0.000E+00 | 0.000E+00 |
| (5,2) | 7.400E-04 | 1.586E-03 |
| (6,1) | 0.000E+00 | 0.000E+00 |
| (6,2) | 0.000E+00 | 0.000E+00 |
| (6,3) | 3.900E-03 | 6.750E-03 |
| (6,4) | 1.480E-02 | 2.352E-02 |
| (7,1) | 0.000E+00 | 0.000E+00 |
| (7,2) | 0.000E+00 | 0.000E+00 |
| (7,3) | 5.000E-04 | 1.305E-03 |
| (7,4) | 2.340E-03 | 4.387E-03 |
| (7,5) | 2.314E-02 | 3.471E-02 |
| (8,1) | 0.000E+00 | 0.000E+00 |
| (8,2) | 0.000E+00 | 0.000E+00 |
| (8,3) | 5.600E-04 | 1.305E-03 |
| (8,4) | 2.180E-03 | 4.265E-03 |
| (8,5) | 1.516E-02 | 2.352E-02 |
| (9,1) | 0.000E+00 | 0.000E+00 |
| (9,2) | 0.000E+00 | 0.000E+00 |
| (9,3) | 1.400E-04 | 3.937E-04 |
| (9,4) | 5.600E-04 | 1.305E-03 |
| (9,5) | 3.460E-03 | 6.228E-03 |
| (10,1) | 0.000E+00 | 0.000E+00 |
| (10,2) | 0.000E+00 | 0.000E+00 |
| (10,3) | 2.000E-05 | 6.429E-05 |
| (10,4) | 6.000E-05 | 1.800E-04 |
| (10,5) | 5.800E-04 | 1.305E-03 |

**Supplementary File 12:**

**Top $10$ regions with significant weighted-mean differences ($q < 0.05$, FDR corrected) in regional variations of local synchrony during the preclinical stages (stages 6 vs 1) [Figure 4D, H, L in the main text]**. The $p$- and $q$-values of 0.000E+00 denote a value less than $1/50,000$, where $50,000$ is the number of bootstrap samplings.

| Frequency band | Regions (AAL3 atlas) | $\delta z$ | $p$-value | $q$-value |
|---|---|---|---|---|
| delta-theta | Right Middle occipital gyrus | 3.089 | 6.000E-05 | 9.246E-05 |
| | Left Middle temporal gyrus | 2.859 | 4.000E-05 | 7.094E-05 |
| | Left Superior temporal gyrus | 2.815 | 2.000E-05 | 4.372E-05 |
| | Right Heschls gyrus | 2.780 | 2.000E-05 | 4.372E-05 |
| | Left Rolandic operculum | 2.767 | 0.000E+00 | 0.000E+00 |
| | Left SupraMarginal gyrus | 2.729 | 4.000E-05 | 7.094E-05 |
| | Left Angular gyrus | 2.726 | 2.000E-05 | 4.372E-05 |
| | Left Heschls gyrus | 2.670 | 0.000E+00 | 0.000E+00 |
| | Left Parahippocampal gyrus | 2.664 | 0.000E+00 | 0.000E+00 |
| | Left Insula | 2.638 | 0.000E+00 | 0.000E+00 |
| alpha | Left Fusiform gyrus | -1.835 | 0.000E+00 | 0.000E+00 |
| | Left Inferior occipital gyrus | -1.700 | 0.000E+00 | 0.000E+00 |
| | Right Fusiform gyrus | -1.671 | 0.000E+00 | 0.000E+00 |
| | Left Lingual gyrus | -1.670 | 0.000E+00 | 0.000E+00 |
| | Left Inferior temporal gyrus | -1.596 | 0.000E+00 | 0.000E+00 |
| | Left Hippocampus | -1.570 | 0.000E+00 | 0.000E+00 |
| | Right Inferior occipital gyrus | -1.554 | 0.000E+00 | 0.000E+00 |
| | Right Parahippocampal gyrus | -1.551 | 0.000E+00 | 0.000E+00 |
| | Left Parahippocampal gyrus | -1.549 | 0.000E+00 | 0.000E+00 |
| | Right Calcarine fissure and surrounding cortex | -1.491 | 0.000E+00 | 0.000E+00 |
| beta | Left Superior temporal gyrus | -2.357 | 0.000E+00 | 0.000E+00 |
| | Left Heschls gyrus | -2.351 | 0.000E+00 | 0.000E+00 |
| | Left Rolandic operculum | -2.258 | 0.000E+00 | 0.000E+00 |
| | Left Angular gyrus | -2.253 | 0.000E+00 | 0.000E+00 |
| | Left Hippocampus | -2.249 | 0.000E+00 | 0.000E+00 |
| | Left Parahippocampal gyrus | -2.227 | 0.000E+00 | 0.000E+00 |
| | Left Insula | -2.227 | 0.000E+00 | 0.000E+00 |
| | Right Heschls gyrus | -2.192 | 0.000E+00 | 0.000E+00 |
| | Left Lenticular nucleus-Putamen | -2.148 | 0.000E+00 | 0.000E+00 |
| | Right Hippocampus | -2.120 | 0.000E+00 | 0.000E+00 |
